# Derivation of the immortalized cell line-UM51-PrePodo-hTERT and its responsiveness to Angiotensin II and activation of RAAS

**DOI:** 10.1101/2022.05.23.493113

**Authors:** Lars Erichsen, Lea Doris Friedel Kloss, Chantelle Thimm, Martina Bohndorf, Kira Schichel, Wasco Wruck, James Adjaye

## Abstract

Recent demographic studies predict there will be a considerable increase of elderly people within the next decades. Aging has been recognized as one of the main risk factors of the world’s most prevalent diseases, including neurodegenerative disorders, cancer, cardiovascular disease, and metabolic disease. During the process of aging a gradual loss of tissue volume and organ function is observed, which is partially caused by replicative senescence. The capacity of cellular proliferation and replicative senescence is tightly regulated by their telomere length. When the telomere length with progressive cell division is critically shortened, the cell becomes proliferative arrested and DNA damage response and cellular senescence are triggered. At this time point the so called “Hayflick limit” is attained.

Podocytes are a cell type that is found in the kidney glomerulus where they have major implications in blood filtration. Mature podocytes are terminal differentiated cells that are unable to undergo cell division *in vivo*. For this reason, the establishment of podocyte cell cultures has been very challenging. In our present study, we present the successful immortalization of a human podocyte progenitor cell line, of which the primary cell cells were isolated directly from the urine of a 51-year-old male. The immortalized cell line has been cultured over the course of one year (∼100 passages) with high proliferation capacity, while still endowed with contact inhibition and P53 expression and activation. Furthermore, by immunofluorescent-based expression and quantitative Real-Time PCR for the podocyte markers NPHS1, SYNPO and WT1 we confirmed the differentiation capacity of the immortalized cells. Finally, we evaluated and confirmed the responsiveness of the immortalized cells on the main mediator Angiotensin II (ANGII) of the renin-angiotensin-system (RAS). Elevated levels of ANGII have been identified as a main risk factor for the initiation and progression of chronic kidney disease (CKD). CKD is characterized by an impairment of podocyte function and subsequently podocyte apoptosis. As a major risk factor for patients to develop CKD, diabetes, hypertension, heart disease and stroke have been recognized - all of which show increased incidence in elderly people. In conclusion, we have shown that it is possible to by-pass cellular replicative senescence (Hayflicks limit) by TERT-driven immortalization of human urine-derived pre-podocyte cells from a 51-year-old African male.

## Introduction

Podocytes are a distinct cell type within the kidney glomerulus. Their major task is the filtration of blood to generate urine and thereby retaining plasma proteins [2]. Therefore, they are part of the glomerular filtration barrier, together with glomerular epithelial cells (GECs) and the glomerular basement membrane (GBM) [3]. To execute this complex task, podocytes need to be terminally differentiated and develop an elaborate and highly specialized actin cytoskeleton. A typical podocyte morphology is composed of three major segments, the cell body, major processes and foot processes (FPs) [4]. Major processes are connected to the cell body of podocytes and can split into foot processes. Neighboring foot processes from different podocytes form an inter-digitating pattern, which is bridged by slit diaphragms (SDs) [4].

During the development of the human kidney, the precursor cells of podocytes arise from SIX2-positive renal progenitor cells. Subsequent developmental processes are then needed to acquire the terminally differentiated state and maturity [4, 5]. Glomerular development proceeds in four stages: the renal vesicle stage, the S-shaped body stage, the capillary loop stage, and the maturing-glomeruli stage [6, 7]. During the S-shaped body stage the podocyte precursors are simply undifferentiated epithelial cells, with apically located tight junctions [8] and prominent mitotic activity [9]. During this specific time point expression of Podocalyxin (PODXL) [10] and the tight junction protein ZO-1 [11] starts, while expression of the podocyte transcription factor WT1 is the highest [12]. When these immature podocytes enter the capillary loop stage, their mitotic activity is abolished and they begin to establish their complex actin cytoskeleton, including the appearance of FPs and the reorganization of cell-cell junctions into SDs [8]. The filtration slits are formed by the spatial formation of the FPs. These are spanned by the glomerular slit diaphragm (SD), which is similar to an adherens junctions structure [13]. For the formation of SDs, the protein Nephrin (NPHS1) is mainly required, which is additionally associated with the actin cytoskeleton and thus contributes to the actin dynamics of podocytes and the formation of FPs [14].

The phenotypical changes occurring between the S-shaped body stage and the capillary loop stage are accompanied by Synaptopodin (SYNPO) [15] and Vimentin [16] expression. Therefore, SYNPO has been recognized as a key marker of differentiated post mitotic podocytes [15, 17, 18], which cannot be detected in undifferentiated or dedifferentiated cells [19, 20].

Numerous kidney diseases, such as chronic kidney disease, membranous nephropathy, congenital nephrotic syndrome and Alport syndrome are associated with proteinuria and or hematuria which is mainly caused by defects in the GBM or alterations in structure and or function of the podocytes itself. Modelling of these pathological conditions has been very challenging in the past. One obvious drawback of studying human podocytopathies is the limited proliferation capacity of these cells. This was further aggravated by the fact that the derivation of kidney originated cells is very challenging. This is particularly true for podocytes, since their complex architecture is not well preserved from kidney biopsy tissue [17]. For these reasons disease modelling of human kidney alterations is often studied in animal models which only replicate human conditions to a certain extend. Therefore, other models have been implemented to study human kidney disease, like the temperature-sensitive SV40 conditionally immortalised podocyte cell line [17] and many iPSC based differentiation protocols for cells of kidney origin [21–25].

Here we describe the successful immortalization of a human podocyte progenitor cell line, obtained directly from urine. The cells show a stable morphology and proliferation capacity *in vitro* for more than one year of cell culture. They express typical podocyte markers, such as NPHS1 and WT1 and upon our recently reported differentiation protocol their proliferation capacity is retained, and they express several podocyte markers such as SYNPO. Due to their high proliferation capacity and their easy handling, our immortalized UM51-hTERT-Pre-Podo cell line offers a unique tool for studying nephrogenesis and kidney-associated diseases.

## Material and methods

### Cell culture conditions

The cell line UM51-PrePodo was derived from the urine of a 51-year-old male of African origin. The cells were cultured, and differentiation was induced as described in [26]. Immortalization of the cells was achieved by lipofection of the pCDNA-3xHA-hTERT plasmid with Xfect (Takara BIO INC, Kusatsu, prefecture Shiga, Japan). The plasmid pCDNA-3xHA-hTERT was obtained from (Addgene plasmid # 51637 ; http://n2t.net/addgene:51637 ; RRID:Addgene_51637) [27]. In brief, 2 µg of the plasmid were incubated at RT for 10 min with 100 µl of transfection buffer and 1 µl of transfection reagent. After incubation, the mix was added to at least 50% confluent 6-well of UM51-PrePodo, growing as a monolayer. In another transfection, 2 µg of the commercial pmaxGFP Vector (Lonza, Swiss, Basel) were incubated at RT for 10 min with 100 µl of transfection buffer and 1 µl of transfection reagent. After incubation, the mix was added to at least 50% confluent 6-well of UM51-PrePodo-hTERT, growing as monolayer.

### Immunofluorescence-based detection of protein expression

Cells were fixed with 4% paraformaldehyde (PFA) (Polysciences, Warrington, United States). Unspecific binding sites were blocked by incubation with blocking buffer, containing 10% normal goat or donkey serum, 1% BSA, 0.5% Triton, and 0.05% Tween, for 2 h at room temperature. Incubation of the primary antibody was performed at 4 °C overnight in staining buffer (blocking buffer diluted 1:1 with PBS). After at least 16 h of incubation, the cells were washed three times with PBS/0.05% Tween and incubated with a 1:500 dilution of secondary antibodies. After three additional washing steps with PBS/0.05% Tween the cell and nuclei were stained with Hoechst 1:5000 (Thermo Fisher Scientific, Waltham, United States). Images were captured using a fluorescence microscope (LSM700; Zeiss, Oberkochen, Germany) with Zenblue software (Zeiss). Individual channel images were processed and merged with Fiji. Detailed Information of the used antibodies are given in supplementary table 1.

### Western Blotting

Protein isolation was performed by lysis of the cells in RIPA buffer (Sigma-Aldrich, St. Louis, United States), supplemented with complete protease and phosphatase inhibtors (Roche, Basel, Switzerland). Proteins were separated on a 7.5% Bis-Tris gel and blotted onto a 0.45 µm nitrocellulose membrane (GE Healthcare Life Sciences, Chalfont St. Giles, United Kingdom). After blocking the membranes with 5% skimmed milk in Tris-buffered Saline Tween (TBS-T), they were incubated overnight with the respective primary antibodies (supplementary table 1). Membranes were washed 3x for 10 min with TBS-T. Secondary antibody incubation was performed for 1h at RT and membranes were subsequently washed 3x for 10 min with TBS-T. Amersham ECL Prime Western Blotting Detection Reagent was used for the chemiluminescent detection (GE Healthcare Life Sciences) and captured with the imaging device Fusion FX.

### Southern Blotting

Genomic DNA-isolation was performed with the DNeasy® Blood & Tissue Kit (Qiagen, Hilden, Germany) according to the manusfactors instructions. 10 µg of the isolated DNA was digested by incubation with EcoRI and HindIII overnight and separated on a 1% agarose gel for 2 h. Prior to blotting of the DNA onto a positively charged nylon membrane (GE Healthcare Life Sciences), the DNA in the gel was denaturated by incubation in 0.25 M HCL solution for 5min followed by 2x/15 min in denaturation solution and 2x/15 min in neutralization solution. DNA transfer was carried out overnight at room temperature with 10x saline-sodium citrate (SSC) buffer. Subsequently the membrane was fixed at 120 °C for 20–35 min and washed with 2x SSC twice. Telomere fragments were detected by a prehybridization step with DIG Easy Hyb Granules for 1h and by the actual hybridization of the membrane with a solution containing Telomere Probe dissolved 1:5000 in DIG Easy Hyb granules (Sigma Aldrich) at 42 °C overnight. Two stringent washing steps were performed after hybridization, blocking was done in blocking solution followed by Anti-DIG-Antibody solution. For detection, the membranes were washed in detection buffer for 5 min and 1 ml substrate solution was added. Pictures were taken with the imaging device Fusion FX after an incubation period of at least 5 min.

### Relative Quantification of podocyte-associated gene expression by real-time PCR

Real time PCR of podocyte-associated gene expression was performed as follows:

Real time measurements were carried out on the Step One Plus Real Time PCR Systems using MicroAmp Fast optical 384 Well Reaction Plate and Power Sybr Green PCR Master Mix (Applied Biosystems, Foster City, United States). The amplification conditions were denaturation at 95 °C for 13 min. followed by 37 cycles of 95 °C for 50 s, 60 °C for 45 s and 72 °C for 30 s. Primer sequences are listed in supplementary table 2.

### Resazurin Assay

Proliferation of UM51-PrePodo cells grown in Proliferation Medium and Advanced RPMI was measured with a resazurin assay (Sigma-Aldrich) according to the manufacturer’s instructions, in brief:

Per condition (PM/adv. RPMI) 50.000 cells were seeded in 500 µl of the respective medium in triplicates. 10x Resazurin solution was mixed with the two media in a 1:10 dilution and cells were incubated at 37 °C and 5% CO2 for 2 h. 200 µl medium for each condition was transferred into a 96-well plate and the absorbance at 570 and 600 nm was measured with the Plate Reader AF2200 (Eppendorf, Hamburg, Germany).

## Results

### Successful immortalization of UM51-PrePodo primary cells

UM51-PrePodo cells were isolated directly from human urine as described in [28]. The cells were transfected with the plasmid pCDNA-3xHA-hTERT [27]. After four months of continual transfections immortalization was achieved and the cells referred to as -UM51-PrePodo-hTERT. This cell line exhibited a much faster proliferation capacity compared to the original primary cells-UM51-PrePodo, and kept proliferating for more than six months, while showing similar morphology (fig. 1a). Both cell lines grow as monolayers, show contact inhibition and a cobblestone-like morphology typical for epithelial cells. We further analyzed P53 expression and phosphorylation by immunofluorescence-based protein detection (fig. 1b). While most of the cells express P53, only a small percentage were positively stained for phosphorylated P53. Since the transfected hTERT protein is N-terminally tagged with 3x HA, successful transfection of the cells was evaluated by Western blotting and immunocytochemistry with a primary antibody recognizing the HA-tag (fig. 1c+e). Generally, the transformed cells tend to appear a little smaller and more elongated than UM51-PrePodo. Since ectopic expression of hTERT has been associated with chromosomal alterations and genomic instabilities [29–32], we analyzed the karyotype of UM51-PrePodo-hTERT at the Institute of Human Genetics at the University Hospital of Düsseldorf (fig. 1d). In total 20 metaphases were analyzed and a hypertriploid karyotype with several chromosomal alterations was observed in all of them.

**Figure 1:**
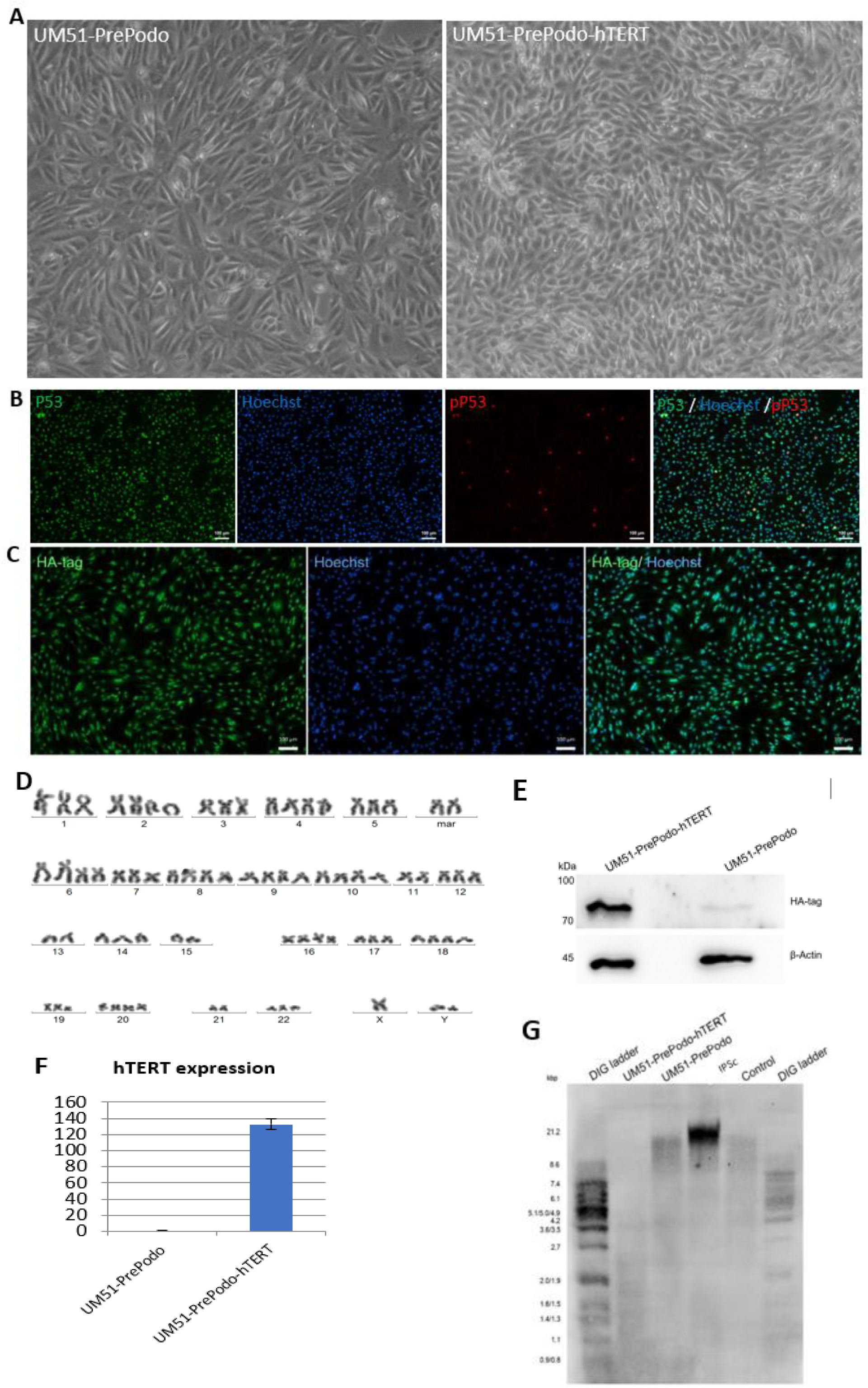
Successful immortalization of the UM51-PrePodo cell line. UM51-PrePodo cell line was successfully immortalized by transfection with the pCDNA-3xHA-hTERT, resembling a similar morphology after transfection as observed by light microscopy (a). Karyotyping of UM51-PrePodo-hTERT revealed a hypertriploid karyotype with several alterations (d). P53 and phosphorylated p53 expression in the immortalized cells was monitored by immunofluorescent based detection (b) (scale bars: 100 µm). Integration of the plasmid was visualized by immunofluorescence-based and Western blot detection of the HA-tag (c+e). Expression of the hTERT gene was determined by quantitative real-time PCR (f). Telomere length was measured by Southern blotting (g).

Karyotype:73-78,X,- ?X/Y,?X/Y,add(1)(q25),+add(1)(q11),del(2)(p11),+del(2)(p11),+del(3)(p13),add(6)(q25),+add(6)(q25),- 9,-11,add(11)(p11),-13,-14,+16,+add(17)(p11),+add(17)(p11),+19,+20,+20,+mar,+mar[cp20].

Since alterations of this magnitude might supposed to have an impact on transcriptional activity, they also might make the cells prone to cancerous transformation. The immunofluorescence-based detection detected expression of the HA-tag in UM51-PrePodo-hTERT, as fluorescence was visible in all cell nuclei. Furthermore, Western blot analysis showed a band at approximately 80 kDa in the protein lysate of UM51-PrePodo-hTERT, which was absent in the non-transfected cell line UM51-PrePodo. ß-Actin was used as loading control and detected at size of 45 kDa (full size western blot images are given in supplementary figure 1). In addition, hTERT gene expression was determined by quantitative real-time PCR, revealing a 132-fold increase of mRNA expression (fig. 1f). Telomere length of UM51-PrePodo-hTERT cells in comparison to the primary UM51-PrePodo cells was evaluated by Southern blotting. Terminal restriction fragments (TRF) containing telomeres and a sub-telomeric region were detected in the control DNA originating from an immortal cell line as supplied by Sigma Aldrich, the induced pluripotent stem cell line A4 as well as the UM51-PrePodo and UM51-PrePodo-hTERT (fig. 1g). TRFs of the A4 cell line were found to be the longest with and length of 13.9 kbp, followed by the control with a fragment size of 7.4 kbp, UM51-PrePodo of 6.3 kbp and UM51-PrePodo-hTERT exhibited to have the shortest TRFs with a size of 1.2 kb.

### Characterization UM51-PrePodo-hTERT

To determine the effect of hTERT driven immortalization of the primary UM51-PrePodo cells on their differentiation capacity, we performed immunofluorescence-based detection of WT1 and NPHS1 protein and real-time PCR based detection of mRNA expression for the genes *WT1, NPHS1, CD2AP, CD24* and *CD106* (fig. 2a-e). As indicated by the immunofluorescence staining the primary UM51-PrePodo cells as well as the immortalized UM51-PrePodo-hTERT cell line both express key podocyte proteins WT1 and NPHS1 (fig. 2 a-d). While real-time PCR based detection revealed a significant increase in mRNA expression for *WT1, NPHS1* and *CD106*, by 0.77 (p=0,001), 2.83 (p=0,03) and 1.66 -fold (p=0,05) respectively, a significant downregulation of *CD2AP* and *CD24*, by 0.57 (p=0,05) and 0.52 (p=0,05) were observed. To test transfection efficacy of the UM51-PrePodo-hTERT cell line, we lipofected freshly plated cells at approximately 60 - 70% confluency with the pmaxGFP Vector and obtained 70% GFP positive cells after 24h (supplementary figure 2).

**Figure 2:**
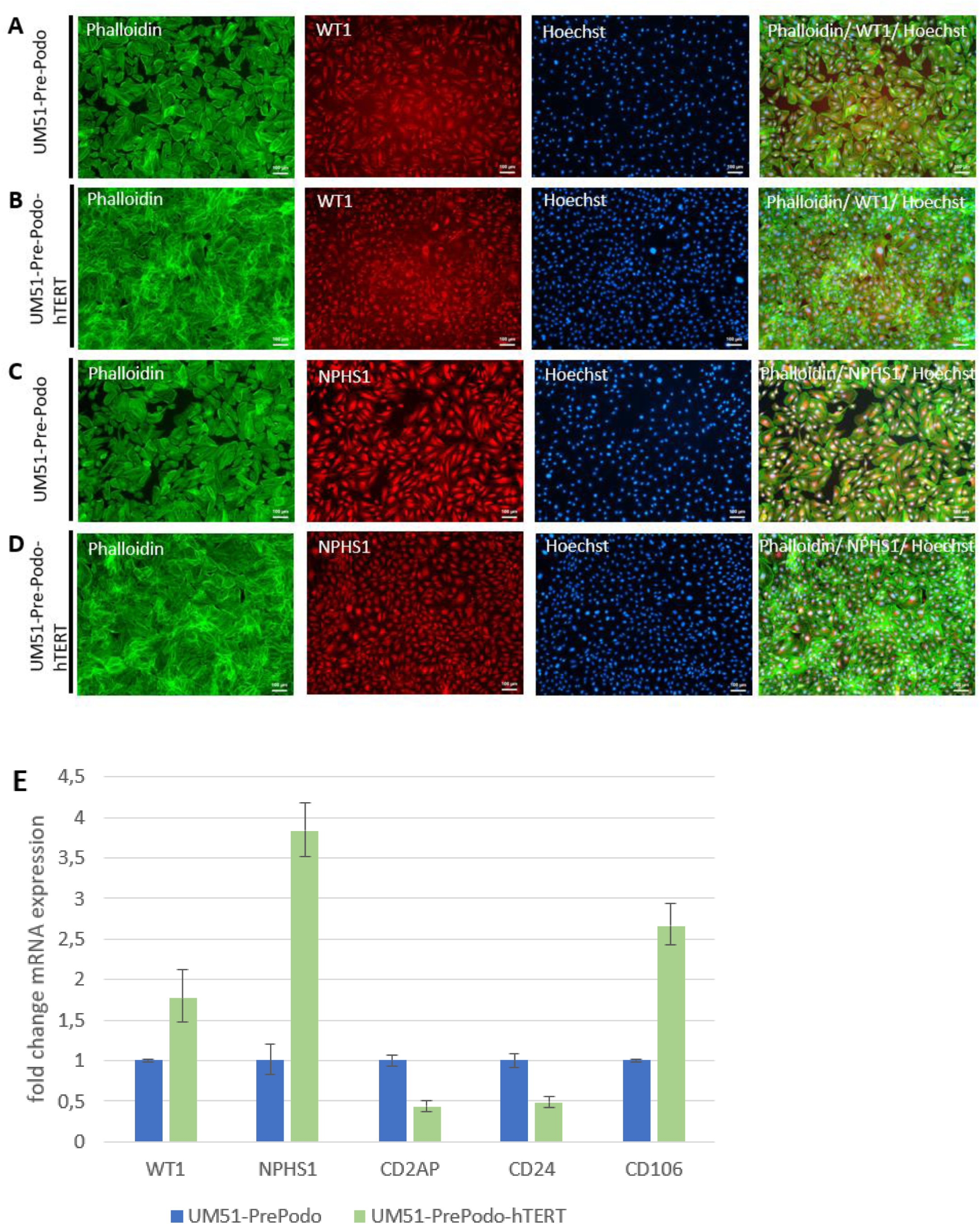
Characterization of immortalized UM51-PrePodo-hTERT. WT1 and NPHS1 expression in the parental UM51-PrePodo as well as the immortalized UM51-PrePodo-hTERT cells was evaluated by immunofluorescence-based detection (a-d) (scale bars: 100 µm). Immunofluorescence-based detection of the parental UM51-PrePodo is given in a and c, while UM51-PrePodo-hTERT is given in b and d. Cytoskeleton was stained with phalloidin. mRNA expression of *WT1, NPHS1, CD2AP, CD24* and *CD106* was determined by quantitative real-time PCR (e).

### Culturing UM51-PrePodo-hTERT in adv. RPMI initiates podocyte differentiation

We recently reported a detailed protocol for the differentiation of SIX2-positive urine derived renal progenitor cells (UdRPCs) into mature podocytes [26]. We applied this protocol to differentiate the UM51-PrePodo-hTERT cell line into podocytes. To determine the proliferation rate of UM51-PrePodo-hTERT cultured under the differentiation conditions, we applied a resazurin assay for 72 h (fig. 3a). The fold-change of normalized absorbance was found to increase in UM51-PrePodo-hTERT cultured in PM medium by 0.8 - fold after 48 h and by additional 0.74 - fold after 72 h. In contrast UM51-PrePodo-hTERT cultured in adv. RPMI medium showed only minor changes in absorbance with 0.03 - fold after 48 h and 0.13 -fold after 72 h. The difference in proliferation rate was also observed using light microscopy (supplementary figure 3). While UM51-PrePodo-hTERT cells cultured in proliferation medium were at 100% confluency after 72 h, cells cultured in advanced RPMI were approximately 60% confluent. Relative protein expression normalized to ß-ACTIN for the HA-tag, for the key podocyte marker SYNPO as well as P53 were detected by Western blotting (fig. 3b). All proteins were detected at the expected sizes of 80 kDa for the HA-tag, 110 kDa for SYNPO, 53 kDa for p53 and 45 kDa for ß- ACTIN. While the most significant difference in protein expression was observed for SYNPO with a 4.8 - fold (p=0,04), HA-tag expression decreased with a fold change of 0.6 (p=0,05). Interestingly the protein level of P53 was unaffected by the culturing medium (full size western blot images are given in supplementary figure 4). Expression of the differentiated podocyte marker SYNPO and the proliferation marker KI-67 were assessed by immunofluorescence-based protein detection (fig. 3c-f). KI-67-protein was detected in the nuclei of UM51-PrePodo-hTERT cells cultured in PM (fig. 3c) as well as adv. RPMI medium (fig. 3d). As expected, a lower percentage of cells grown in adv. RPMI expressed the proliferation marker than cells cultured in PM-3.8% compared to 13.9% respectively (fig. 3g). While UM51-PrePodo-hTERT cells grown in PM lacked SYNPO protein expression (fig. 3e), a clear signal could be detected in cells grown in adv. RPMI (fig. 3f). Additionally, we measured the expression of *CD24, CD106, SYNPO, P53* and *KI67* by quantitative real-time PCR (fig. 3h). In accordance with the Western blot and immunofluorescence-based protein detection, the mRNA level of *SYNPO* was found to be significantly increased (3.2 - fold; p=0,01), while *P53* and *KI67* were found to be downregulated (0.41, p=0,01 and 0.82 -fold, p=0,01 respectively). Furthermore, mRNA expression of the surface marker CD106 was found to be significantly downregulated (0.74 - fold, p=0,04) and CD24 expression was found to be unaltered.

**Figure 3:**
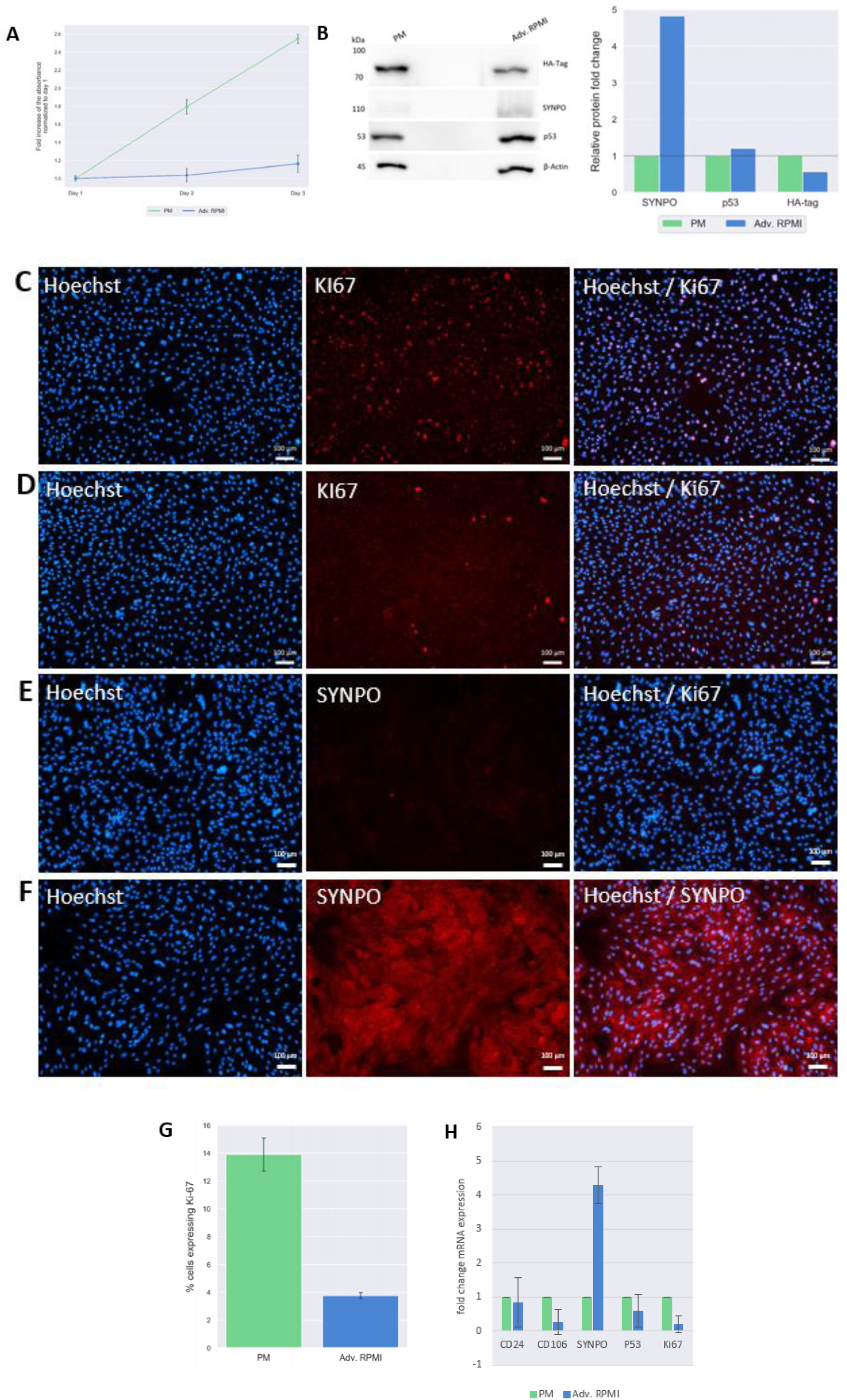
Culturing UM51-PrePodo-hTERT in adv. RPMI initiates podocyte differentiation. Proliferation of immortalized UM51-PrePodo-hTERT cells cultured in PM and adv. RPMI medium was assessed by a resazurin assay (a). Relative protein expression normalized to ß-ACTIN for the HA-tag, SYNPO and P53 was detected by Western blot (b). Ki67 and SYNPO expression in UM51-PrePodo-hTERT cultured in PM and adv. RPMI was detected by immunofluorescence-based staining (c-f).Immunofluorescence-t based detection of UM51-PrePodo-hTERT cells grown in PM are given in c and e, while cells grown in adv. RPMI are given in d and f (scale bars: 100 µm). Percentage of UM51-PrePodo-hTERT cells cultured in Proliferation medium and Advanced RPMI expressing the proliferation marker Ki-67 is given in g. mRNA expression of *CD24, CD106, SYNPO, P53 and KI67* was determined by quantitative real time PCR (h).

### Comparative transcriptome analysis of Urine derived Renal Progenitor cells UM51 with the immortalized UM51-PrePodo-hTERT cell line

After the successful immortalization of UM51-PrePodo into UM51-PrePodo-hTERT and differentiation with adv. RPMI we performed a comparative transcriptome analysis. Hierarchical clustering analysis comparing the transcriptomes of UM51-PrePodo and UM51 podocytes with their respective immortalized counterpart revealed a distinct expression pattern of wild type and immortalized cells (fig. 4a). By comparing the expressed genes (det-p < 0.05) 582 are exclusively expressed in the UM51-PrePodo and 703 in UM51-PrePodo-hTERT grown in PM (fig. 4b). In total 13914 genes were found to be expressed in common between UM51-PrePodo and UM51-PrePodo-hTERT grown in PM (fig. 4b). After differentiation of UM51-PrePodo and UM51-PrePodo-hTERT by culturing the cells in adv RPMI supplemented with 30 µM retinoic acid, transcriptome data revealed 880 genes to be exclusively expressed in UM51 podocytes and 460 in UM51-PrePodo-hTERT grown in adv RPMI (fig 4c). In total 13922 genes were found to be expressed in common between UM51-PrePodo-hTERT grown in adv. RPMI and UM51-PrePodo-hTERT grown in PM (fig. 4d). In total 13699 genes were found to be expressed in common between UM51-Podocytes and UM51-PrePodo-hTERT grown in adv. RPMI (fig. 4c). By comparing the expressed genes (det-p < 0.05) 237 are exclusively expressed in the UM51-PrePodo-hTERT grown in adv. RPMI and 695 in UM51-PrePodo-hTERT grown in PM (fig. 4d). Wild-type cells and their immortalized counterparts share many of the most over-represented GO BP-terms, such as regulation of ion transport, regulation of organic acid transport and developmental processes. The most over-represented GO BP-terms exclusive to UM51-PrePodo are associated with planar cell polarity involved in axis elongation and skin development. In comparison the most over-represented GO BP-terms exclusive expressed in the immortalized UM51-PrePodo-hTERT cells grown in PM are associated with cell cycle checkpoints, nuclear division, regulation of mitotic cycle and DNA replication (fig. 4e). The most over-represented GO BP-terms exclusive to UM51 podocytes are associated with regulation of ion transport, heterophilic cell-cell adhesion via plasma membrane cell adhesion molecules and regulation of system processes. In comparison the most over-represented GO BP-terms exclusive expressed in the immortalized UM51-PrePodo-hTERT cells grown in adv RPMI are associated with chromosome segregation, cell cycle checkpoints, DNA replication, Cell cycle and regulation of DNA replication (fig. 4f). UM51-PrePodo-hTERT cells grown in both media share many of the most over-represented GO BP-terms, such as cell junction organization, urogenital system development, potassium ion transport and GPCR ligand binding. The most over-represented GO BP-terms exclusive to UM51-PrePodo-hTERT grown in adv. RPMI are associated with specification of animal organ identity and chemotaxis. In comparison the most over-represented GO BP-terms exclusive expressed in the immortalized UM51-PrePodo-hTERT cells grown in PM are associated with embryonic pattern specification, embryonic organ development and positive regulation of cell-cell adhesion (fig. 4g). Additionally, the transcriptomes of UM51-PrePodo, their derived podocytes and the immortalized UM51-PrePodo-hTERT grown in PM and adv. RPMI were compared with expressed genes in iPS cell– derived podocytes, kidney biopsy isolated human glomeruli, and mouse podocytes (fig. 4h), as reported by Sharmin et al. [33]. This analysis revealed that many key genes encoding for podocyte function and structure are upregulated upon differentiation with adv. RMPI in both UM51-PrePodo and UM51-PrePodo-hTERT cells, such as *SYNPO, SEMA3G, NPHS1, NPHS2, LAMC1, INF2, SPARC, FYN* and *PODXL*. While the expression of some of the genes listed by Sharmin et al. were found to be highly enriched in UM51-PrePodo-hTERT cells cultured in both media, such as *NLK, TCF21, CPD, SGMS1, PLOD2, IL13RA1, GRK5, RAPBP, HTRA1, PTPRO, WT1, NES* and *DDN*, the expression of other genes was only induced in the wild-type cells cultured in adv. RPMI, such as *ANXA1, TJP1, PDLIM5, CORO2B, TAGLN2, ELF4* and *AMIGO2*.

**Figure 4:**
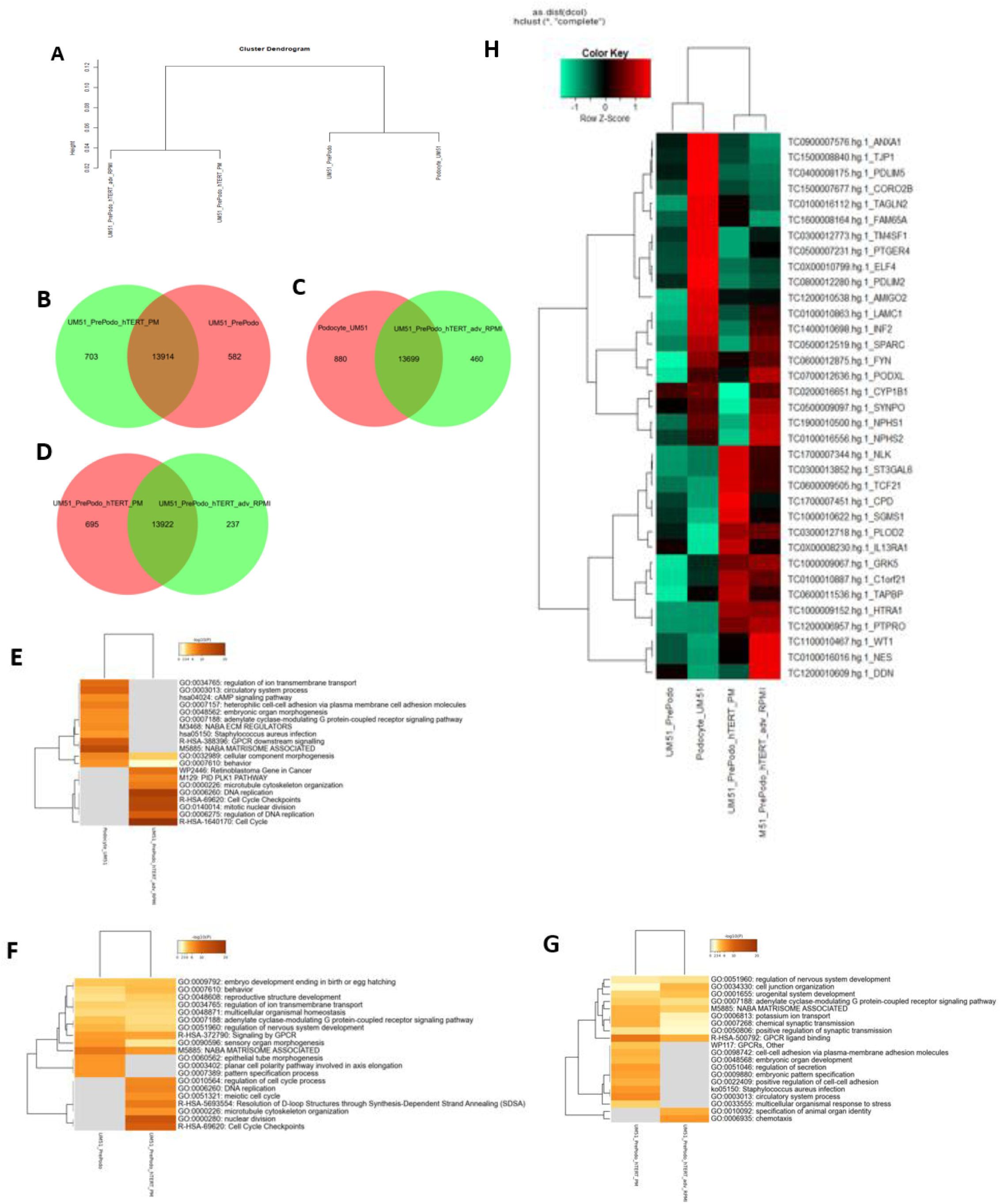
Comparative transcriptome and Gene Ontology analysis of urine-derived renal progenitor UM51 and derived immortalized UM51-PrePodo-hTERT. A hierarchical cluster dendrogram revealed distinct clusters of immortalized and wild type cells (A). Expressed genes (det-p < 0.05) in UdRPCs and podocytes compared in the Venn diagrams (B - D), shows distinct (582 in UM51-PrePodo; 703 in UM51-PrePodo-hTERT in PM, 880 Podocyte UM51; 460 UM51-PrePodo-hTERT in adv RPMI and 605 in UM51-PrePodo-hTERT in PM, 237 UM51-PrePodo- hTERT in adv RPMI) and overlapping (13914, 13699 and 13922) gene expression patterns. The most over-represented GO BP-terms exclusive in either UM51-PrePodo or UM51-PrePodo-hTERT in PM are shown in e. The most over-represented GO BP-terms exclusive in either Podocytes UM51 or UM51-PrePodo-hTERT in adv. RPMI are shown in f. The most over-represented GO BP-terms exclusive in either UM51-PrePodo-hTERT in PM or UM51-PrePodo-hTERT in adv. RPMI are shown in g. Figure h shows a heatmap comparing UM51-PrePod and UM51-PrePodo-hTERT in PM with their differentiated counterparts Podocytes UM51 and UM51-PrePodo-hTERT in adv. RPMI for a geneset commonly expressed in iPS cell–derived podocytes, kidney biopsy isolated human glomeruli, and mouse podocytes.

### Effects of Angiotensin II on the differentiated UM51-Pre-Podo-hTERT podocytes

We recently showed that Angiotensin II (ANGII) treatment in UdRPC differentiated podocytes trigger disruption of the cytoskeleton, resulting in the inhibition of podocyte spreading and subsequent loss of foot processes as observed by a round and condensed morphology [26]. To evaluate the effect of ANGII on the differentiated podocytes derived from the UM51-Pre-Podo-hTERT cell line, we treated the cells with 100 µM ANGII for 24 h and with a combination of 100 µM ANGII and 1 µM of the selective, competitive angiotensin II receptor type 1 antagonist Losartan for 24 h. After 24 h, dynamic changes in morphology and a significant down-regulation of α–ACTININ expression could be observed in cells treated with 100 µM ANGII by immunofluorescence-based detection (fig. 5a). While all untreated podocytes stained positive for α–ACTININ, only a small number of cells appeared to express α–ACTININ after 24 h of ANGII treatment. Furthermore, the control podocytes show the typical “fried egg” podocyte morphology, whereas in ANG II treated podocytes a more rounded and condensed morphology could be observed. To confirm that the disruptive effect is indeed mediated by ANGII, the effects on the cytoskeleton were evaluated by gene-specific mRNA expression of ANGII receptors, *AGTR1* and *AGTR2*, as well as *SNYPO* and *NPHS1* (fig. 5b-e). All genes were found to be significantly upregulated (p<0,05) upon 100 µM ANGII for 24 h except *NPHS1*, which was found to be significantly downregulated (p=0,03). These finding indicates that the effect is mediated via both receptors. ANGII treatment upregulated *AGTR1* expression by 0.4 - fold (fig. 5b) and *AGTR2* by 0.8 - fold (fig. 5c). Interestingly *SNYPO* was also found to be upregulated by 0.5-fold upon ANGII treatment (fig. 5d) while *NPHS1* was found to be downregulated by 0,4 - fold. When cells were treated with the combination of 100 µM ANGII and 1 µM Losartan the expression of all genes were significantly (p<0,05) upregulated. While the expression of *AGTR2, NPHS1* and *SYNPO* was found to be increased by 0,7-, 1.4- and 0,8– fold. Interestingly, mRNA expression of *AGTR1* was significantly upregulated by 24- fold. Furthermore, the Losartan treatment rescued the ANGII-induced disruption of the cytoskeleton as indicated by immunofluorescence-based detection of α–ACTININ expression (fig. 5a). Additionally, the significant upregulation of SYNPO (p=0,03) for both treatments was also confirmed by Western blotting(fig.5e).

**Figure 5:**
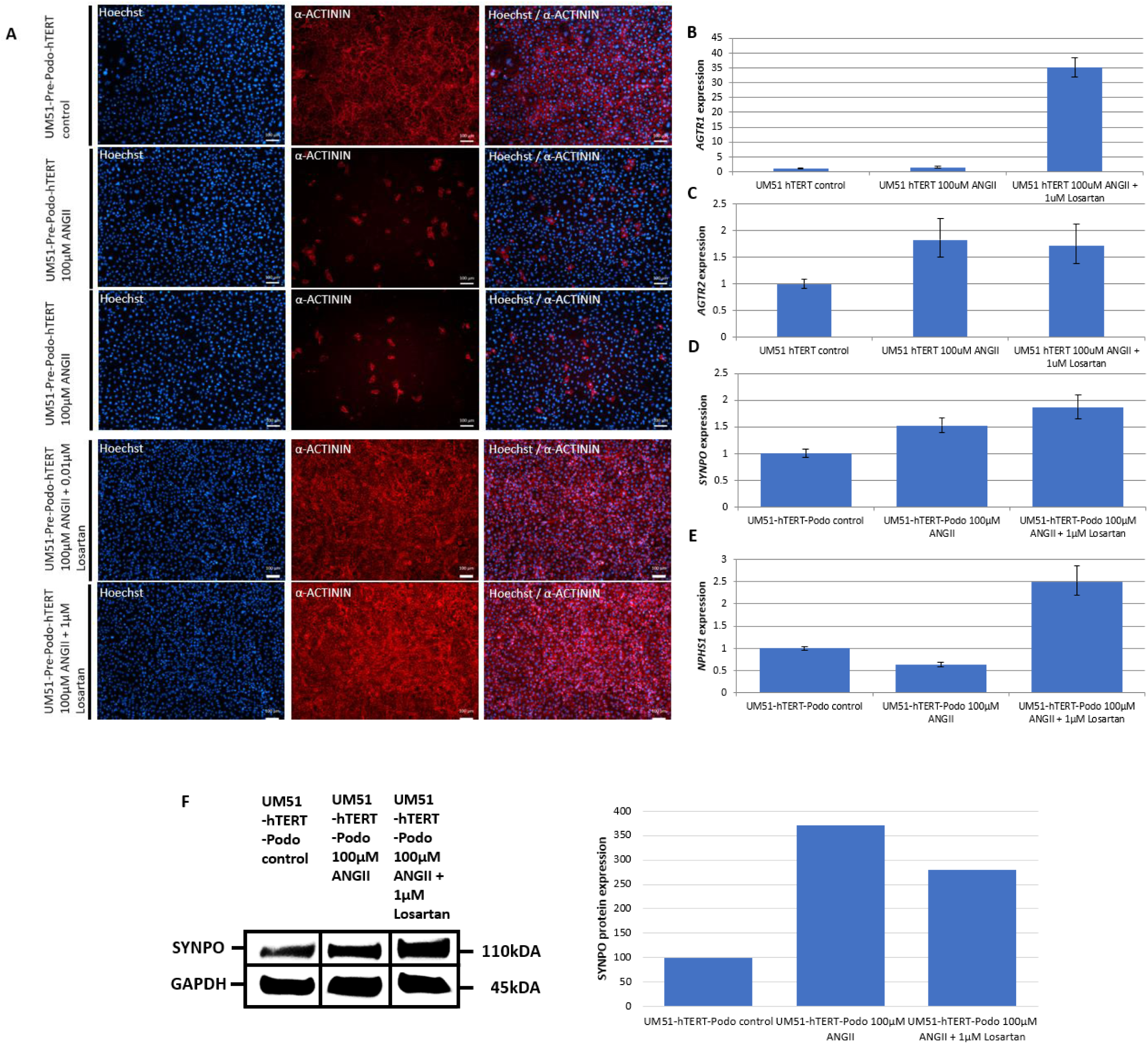
Effects of Angiotensin II on the differentiated UM51-Pre-Podo-hTERT cultured in adv. RPMI. UM51-Pre-Podo-hTERT cultured in adv. RPMI supplemented with 30μM retinoic acid were treated with 100µM ANGII for 24h or a combination of 48h 1µM Losartan and 24h 100µM ANGII. The top panel shows the control morphology. The next two panels show morphology changes after 24h of 100 µM ANGII treatment and the last two panels show the morphology for the combination of Losartan and ANGII. Cytoskeleton was visualized by immunofluorescence-based detection of α –ACTININ in red (a) (scale bars: 50 µm). Expression of ANGII receptors *AGTR1* (b), *AGTR2* (c) and expression of podocyte marker *SYNPO* (d) and NPHS1 (e) were determined by quantitative real time PCR normalized with the ribosomal encoding gene-RPL0. Expression of the podocyte marker Synaptopodin was determined by Western blotting (f).

## Discussion

Podocytes are a cell type that is found in the kidney glomerulus where they have major implications in blood filtration. Since mature podocytes are terminal differentiated cells that are unable to undergo cell division *in vivo [13]*, establishing podocyte cell cultures has been very challenging. In the present manuscript we describe the successful immortalization of a human podocyte progenitor cell line-UM51-PrePodo-hTERT-the primary cells were isolated directly from the urine of a male of African origin. UM51-PrePodo-hTERT has been cultured over the course of one year (∼100 passages) with high proliferation capacity. Numerous cell lines have been immortalized by overexpression of hTERT, leading to chromosomal aneuploidies and alterations in gene expression [34, 35]. Even though UM51-PrePodo-hTERT has a hypotriploid karyotype, the cells do show contact inhibition and P53 expression and activation, indicating that the cells have not lost cell-cycle-checkpoints or undergone cancerous transformation.

The comparison of the transcriptomes pertaining to podocytes differentiated from UM51-PrePodo-hTERT with previously reported data-sets of iPSC–derived podocytes, biopsy derived human glomeruli, and mouse podocytes [33], revealed distinct clustering of UdRPCs and their differentiated counterparts, whereas most of the genes were found to be upregulated after our podocyte differentiation protocol. These results confirm the differentiation capacity of UM51-PrePodo-hTERT into mature podocytes-based on both protein and mRNA expression of NPHS1, SYNPO and WT1. Interestingly, by employing our recently published podocyte differentiation protocol [26], the proliferation capacity of UM51-PrePodo-hTERT was diminished and a significant upregulation of SYNPO expression was observed. SYNPO has been recognized as a key marker of differentiated post-mitotic podocytes [15, 17, 18], which cannot be detected in undifferentiated or dedifferentiated cells [19, 20]. Furthermore, cells in the glomerulus expressing CD133, CD24 and CD166 have been recognized to be highly proliferative, while cells expressing only CD133 and CD24 show lower proliferative capacity and a committed phenotype towards differentiation [36]. Taken together these two findings, expression of SYNPO and loss of CD24 by cultivation of the UM51-PrePodo-hTert line in adv. RPMI and RA confirms the differentiation potential of this cell line into Podocytes.

We propose that UM51-PrePodo-hTERT can a) be differentiated into mature podocytes and b) exit the hTERT-driven cell-cycle progression and enter the G0 phase. Of note, during nephron development podocytes lose their mitotic activity [4] and SYNPO expression is first detectable during the capillary loop stage [15]. This transition is reflected by our KI-67 and SYNPO staining’s in the UM51-hTERT-Pre-Podo cells cultured in adv RPMI.

Finally, we evaluated the responsiveness of UM51-PrePodo-hTERT to Angiotensin II (ANGII) which is a key mediator of the renin-angiotensin-system (RAS) [1, 26]. Elevated levels of ANGII have been identified as a main risk factor for the initiation and progression of chronic kidney disease (CKD). Increased ANGII concentrations are associated with the downregulation of Nephrin and Synaptopodin expression in podocytes [37, 38], leading to podocyte injury [39] and subsequent apoptosis [40]. We observed a disruptive effect of ANGII on the cytoskeleton of podocytes derived from UM51-PrePodo-hTERT, which was accompanied by upregulation of the two Angiotensin II receptors, AGTR1 and AGTR2. Here, it is interesting to note that most cells completely down-regulated α-ACTININ expression, while SYNPO expression was found to be upregulated. This might be explained by the duration of the differentiation. The cells were cultured in adv RPMI for seven days, in which not all cells entered the quiescent state, as indicated by our KI-67 staining and proliferation assay. This makes it tempting to speculate that the hTERT driven immortalization keeps a subpopulation of the immortalized cells in a proliferative state, which then triggered by the culture condition, enables the podocyte lineage specific gene expression repertoire.

There are currently numerous angiotensin receptor blockers (ARBs) and angiotensin-converting enzyme inhibitors (ACEIs) in clinical use for the treatment of a variety of renal diseases [41, 42]. A universal ARB is Losartan, an anti-hypertensive agent from the Sartan group. Losartan is used for the treatment of hypertension, as well as for other therapeutic indications. Its action is based on reversing the effects of ANG II on AGTR1 by the selective masking of AGTR1. Our data provide evidence that the addition of Losartan can counteract the downregulation of the slit diaphragm protein NPHS1 induced by ANG II thus leading to the restoration of podocyte architecture by the upregulated expression of Nephrin. From this it can be deduced that Losartan can efficiently block AGTR1, whereby ANG II binds with a higher affinity presumably to AGTR2. The activation of AGTR2 has has been linked to vasodilation, development, cell differentiation, tissue repair and apoptosis [43, 44]. So, the blockage of AGTR1 and thereby the hypothesized hyperactivation of AGTR2 might restore the expression levels of podocyte-specific genes back to the differentiated healthy state.

To surmise, we have described the successful immortalization of the very first male African Pre-Podocyte cell line-UM51-Pre-Podo-hTERT, which grow as a monolayer, show contact inhibition and a cobblestone-like morphology typical for epithelial cells. UM51-Pre-Podo-hTERT can be (a) efficiently transfected as shown by GFP-encoding vector expression and (b) differentiated into mature podocytes thus amenable for studying nephrogenesis and associated diseases thus obviating the need of iPSCs. Furthermore, the responsiveness of the derived podocytes to ANGII implies potential applications in kidney-associated disease modelling, nephrotoxicity studies and drug screening.

## Supporting information

Table 3 UM51_PrePodo_vs_UM51_PrePodo_hTERT

Table 4 UM51_Podo_vs UM51_PrePodo_hTERT_adv_RPMI

## Informed Consent Statement

In this study, urine samples were collected with the informed consent of the donors.

## Data Availability Statement

Microarray data generated for this study are available at NCBI GEO under the accession number xxx.

## Acknowledgments

James Adjaye acknowledges funding from the medical faculty of Heinrich Heine University, Duesseldorf, Germany.

