## supplement for "Derivation of the immortalized cell line-UM51-PrePodo-hTERT and its responsiveness to Angiotensin II and activation of RAAS"

**Supplemental Material Table of Contents**

Table1: antibodies

Table 2: RT-qPCR Primers

Table 3: UM51_PrePodo_vs_UM51_PrePodo_hTERT

Table 4: UM51_Podo_vs UM51_PrePodo_hTERT_adv_RPMI

Supplementary figure 1: Complete western blot images for the detection of the HA-tag in the cell lines UM51- PrePodo-hTERT and UM51-PrePodo. (A) HA-tag, (B) β-Actin.

Supplementary figure 2: Transfection of UM51-PrePodo-hTERT with the TurboGFP- vector.

Supplementary figure 3: Morphology of UM51-PrePodo-hTERT cells cultured in Proliferation Medium (right) and Advanced RPMI (left) during the resazurin assay.

Supplementary figure 4: Complete western blot images for the comparison of UM51-PrePodo-hTERT cultured in Proliferation Medium (PM) and Advanced RPMI (Adv. RPMI). (A) HA-tag, (B) SYNPO, (C) P53 and (D) β-Actin.

Table1: antibodies

| Antigen | Company | Dilution | |
|---|---|---|---|
| | | ICC | WB |
| α-ACTININ | Abcam (#108198) | 1:200 | / |
| ß-ACTIN | Cell Signaling Technology (3700S) | / | 1:4000 |
| HA-tag | Proteintech (51064-2-AP) | 1:200 | 1:4000 |
| KI67 | Cell Signaling Technology (9449S) | 1:200 | / |
| NPHS1 | Invitrogen (#PA5-20330) | 1:200 | 1:1000 |
| P53 | Merck (OP43-100UG) | 1:200 | 1:1000 |
| pP53 | Cell Signaling Technology (9283S) | 1:200 | / |
| SYNPO | Thermo Fisher Scientific (PA5-56997) | 1:200 | 1:1000 |
| WT1 | Merck (3012219) | 1:200 | / |
| Anti-mouse HRP-labeled | Thermo Fisher Scientific #NA931 | / | 1:4000 |
| Anti-rabbit HRP-labeled | Cell Signaling #7074S | / | 1:1000 |

Table 1: RT-qPCR Primers

| Primer name | Sequence | Annealing temperature (°C) | Product length (bp) |
|---|---|---|---|
| AGTR1 s | 5'- tct cag cat tga tcg ata cc -3' | 60 | 80 |
| AGTR1 as | 5'- tga ctt tgg cta caa gca tt -3' | | |
| AGTR2 s | 5'- tat ggc ctg ttt gtc ctc at -3' | 60 | 114 |
| AGTR2 as | 5'- cat tgg gca tat ttc tca gg -3' | | |
| CD106 s | 5'- cga acc caa aca aag gca gag ta -3' | 60 | 84 |
| CD106 as | 5'- gag gaa ggg ctg acc aag acg -3' | | |
| CD24 s | 5'- gcg gac ttt tct ttt ggg ggg -3' | 60 | 174 |
| CD24 as | 5'- cca gca gca gcc cca g -3' | | |
| CD2AP s | 5'- aac tca tga agc cca gga cga -3' | 60 | 100 |
| CD2AP as | 5'- ctg atc cag atg cag ttt cac tca c -3' | | |
| hTERT s | 5'- cgg aag agt gtc tgg agc aa -3' | 60 | 145 |
| hTERT as | 5'- gga tga agc gga gtc tgg a-3' | | |
| KI67 s | 5'- tcg tcc cag tgg aag agt tg -3' | 60 | 144 |
| KI67 as | 5'- cag ccc cgc tcc ttt tga t -3' | | |
| NPHS1 s | 5'- gcg ggt tct gct acg atg gtg -3' | 60 | 295 |
| NPHS1 as | 5'- caa aca cac cag cct cac ccg -3' | | |
| P53 s | 5'- cag ggc agc tac ggt ttc c-3' | 60 | 102 |
| P53 as | 5'- cag ttg gca aaa cat ctt gtt gag-3' | | |
| RPL s | 5'- tcg aca atg gca gca tct ac -3' | 60 | 195 |
| RPL as | 5'- atc cgt ctc cac aga caa gg -3' | | |
| SYNPO s | 5-' ccc caa cct ctc ctc taa cc -3' | 60 | 116 |
| SYNPO as | 5'- atg aca cag gag gca gaa gaa t -3' | | |
| WT1 s | 5'- cac agc aca ggg tac gag a -3' | 60 | 133 |
| WT1 as | 5'- caa gag tcg ggg cta ctc c -3' | | |

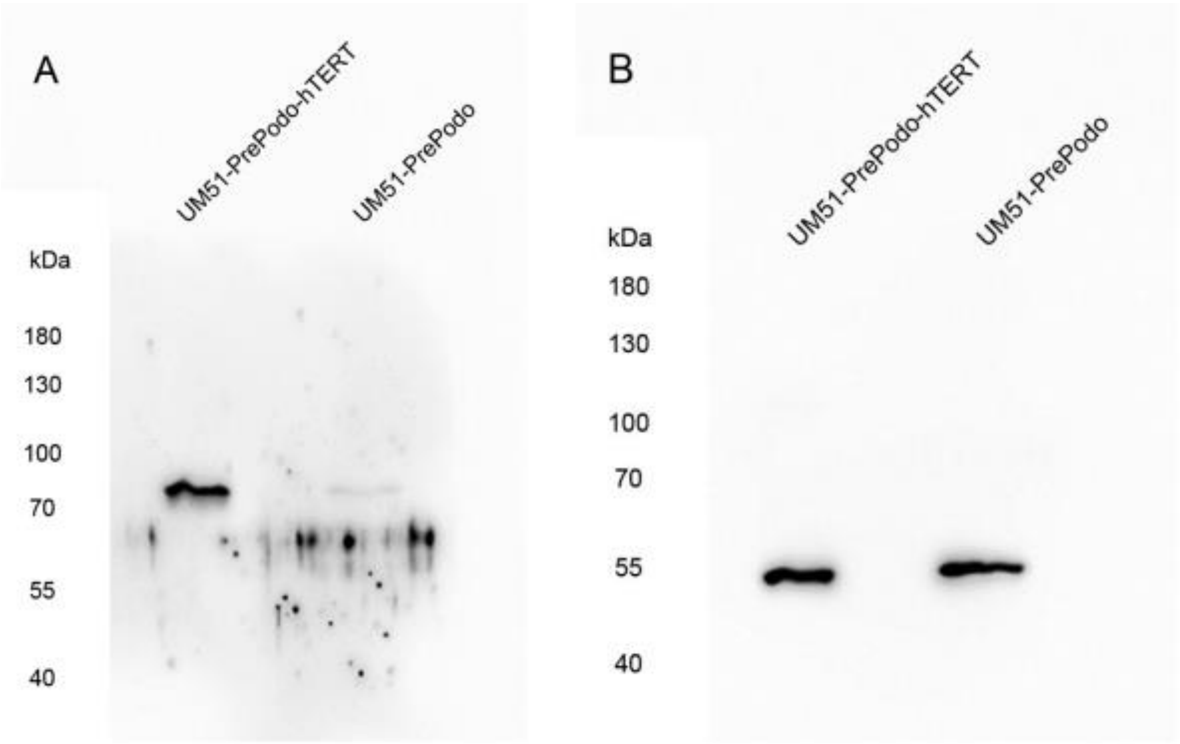


**Supplementary figure 1: Complete western blot images for the detection of the HA-tag in the cell lines UM51- PrePodo-hTERT and UM51-PrePodo. (A) HA-tag, (B) β-Actin.**

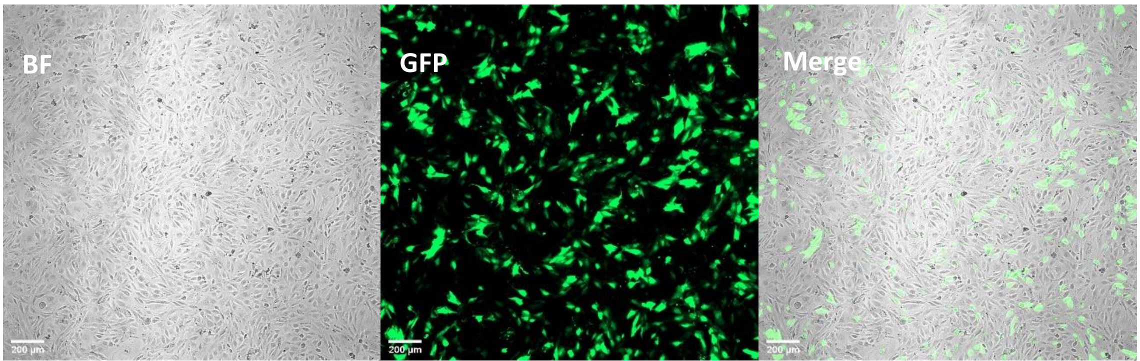

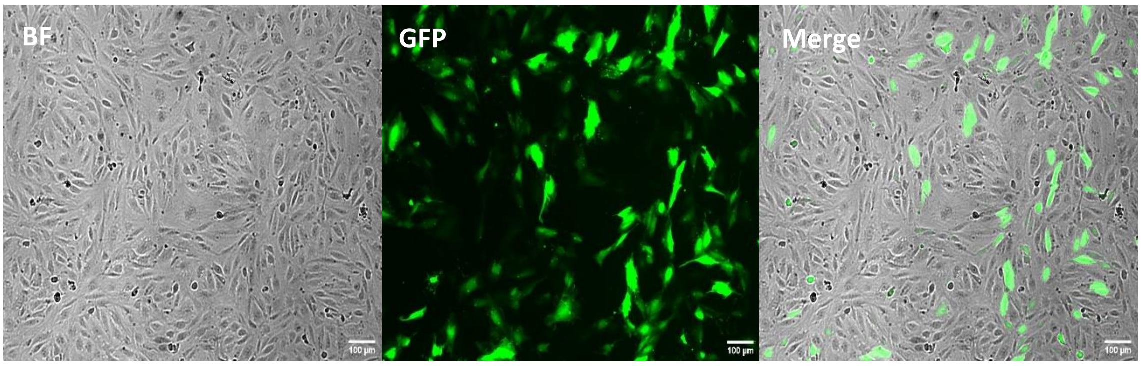


**Supplementary figure 2: Transfection of UM51-PrePodo-hTERT with the TurboGFP- vector.**

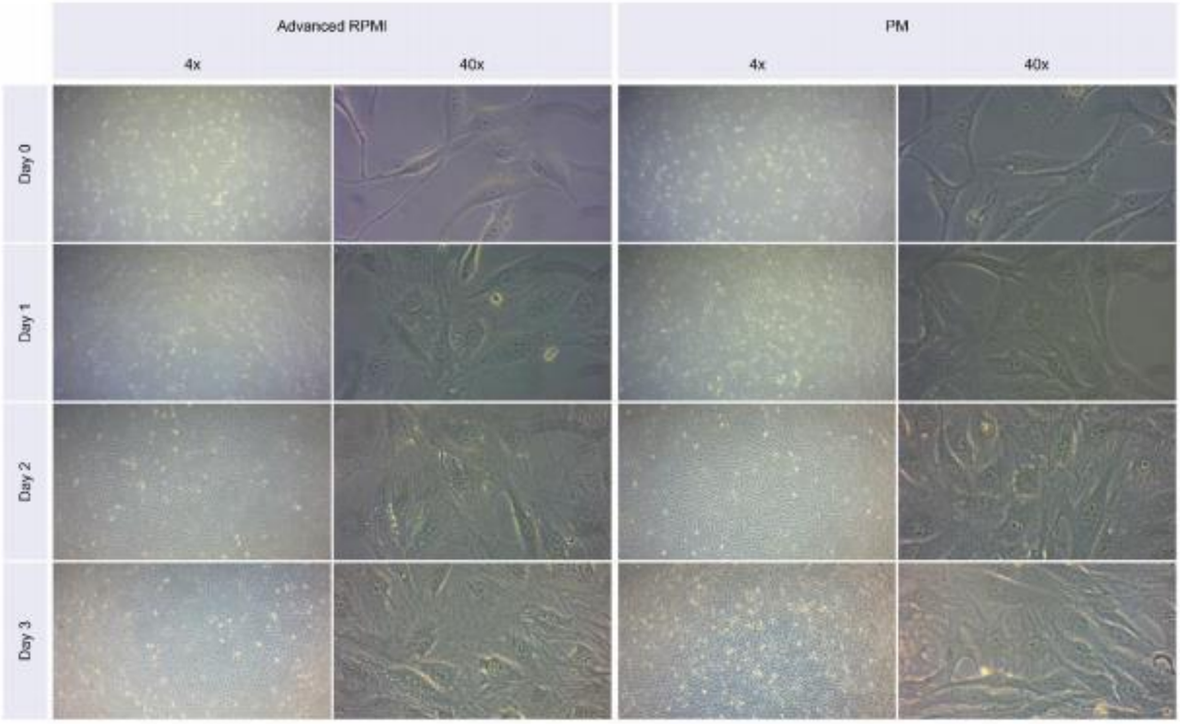


**Supplementary figure 3: Morphology of UM51-PrePodo-hTERT cells cultured in Proliferation Medium (right) and Advanced RPMI (left) during the resazurin assay.**

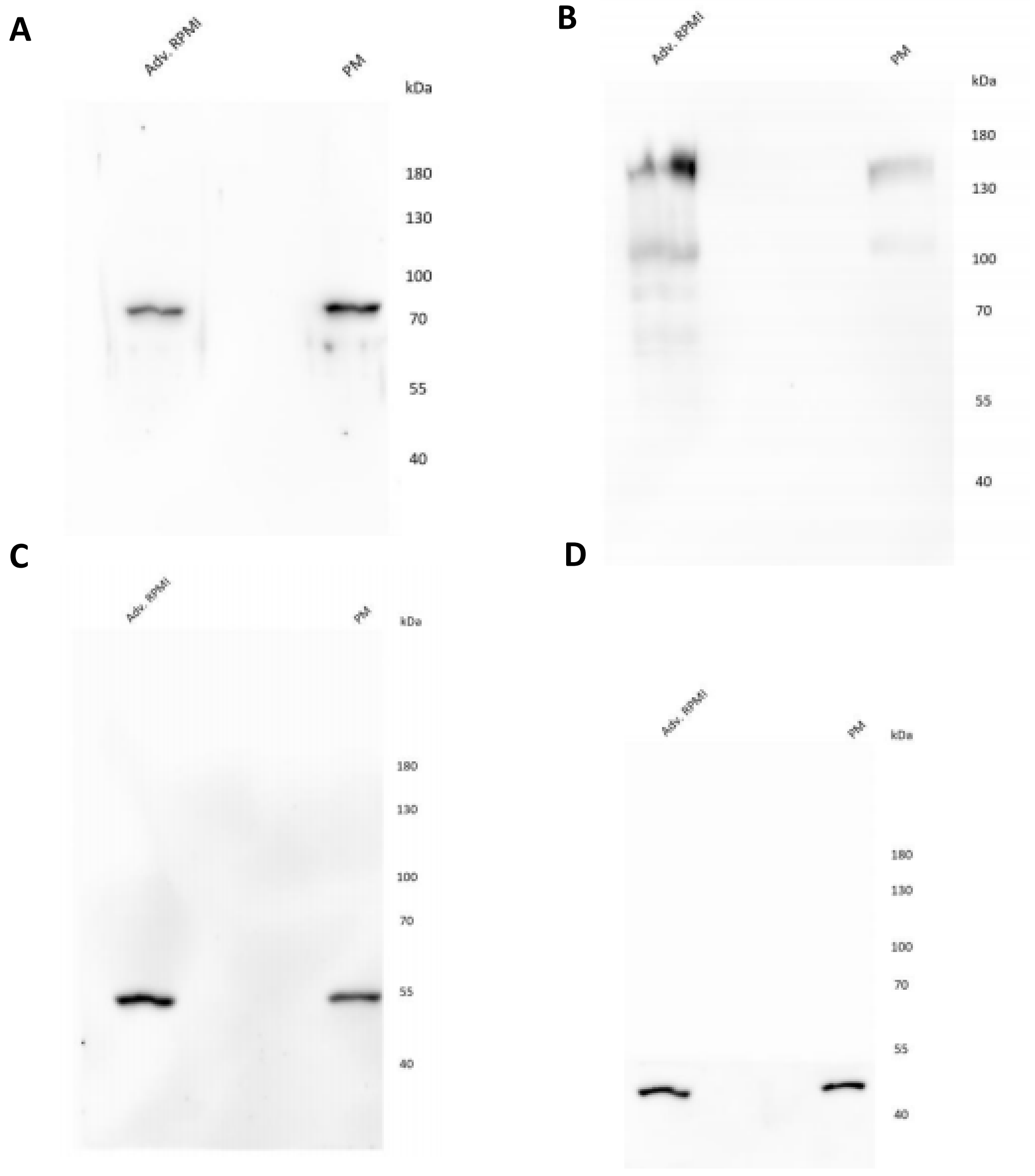


**Supplementary figure 4: Complete western blot images for the comparison of UM51-PrePodo-hTERT cultured in Proliferation Medium (PM) and Advanced RPMI (Adv. RPMI). (A) HA-tag, (B) SYNPO, (C) P53 and (D) β- Actin.**
